# Enteropathogenic *Escherichia coli* induces *Entamoeba histolytica* Lévy-like movement on fibronectin-rich substrate by reducing traction forces

**DOI:** 10.1101/2024.09.27.615376

**Authors:** Yuanning Guo, Jun Ye, Anas Odeh, Meirav Trebicz-Geffen, Haguy Wolfenson, Serge Ankri

**Author notes:** Corresponding author (H.W.), (S.A.). These authors contributed equally to this work (Yuanning Guo, Jun Ye).

## Abstract

Amebiasis, caused by *Entamoeba histolytica*, is a global health concern, affecting millions and causing significant mortality, particularly in areas with poor sanitation. Although recent studies have examined *E. histolytica*’s interaction with human intestinal microbes, the impact of bacterial presence on the parasite’s motility, mechanical forces, and their potential role in altering invasiveness have not been fully elucidated. In this study, we utilized a micropillar-array system combined with live imaging to investigate the effects of enteropathogenic *Escherichia coli* on *E. histolytica*’s motility characteristics, F-actin spatial localization, and traction force exerted on fibronectin-coated substrates. Our findings indicate that co-incubation with *E. coli* significantly enhances the motility of *E. histolytica*, as evidenced by the enhancement of Lévy-like movement patterns, i.e., increased directionality and velocity. This increased motility is accompanied by a reduction in F-actin-dependent traction forces and podosome-like structures on fibronectin-coated substrates, but with increased F-actin localization in the upper part of the cytoplasm. These findings highlight the role of physical interactions and cellular behaviors in modulating the parasite’s virulence, providing new insights into the mechanistic basis of its pathogenicity.

**Author Summary:** Amebiasis, caused by the protozoan parasite *Entamoeba histolytica*, is a major global health issue, affecting around 50 million people and resulting in 100,000 deaths annually. The disease is transmitted through contaminated food and water. Upon reaching the intestines, which are teeming with bacteria, *E. histolytica* begins its invasion by removing the protective mucus layer, followed by adhering to and detaching enterocytes, leading to the disruption and degradation of the epithelial barrier. Afterwards, *E. histolytica* invades along the fibronectin-rich basement membrane deep into the crypts of Lieberkühn, eventually penetrating the fibronectin-rich lamina propria. This leads to further tissue destruction and potential dissemination to distant organs, causing severe complications. In our study, we explored how the presence of enteropathogenic bacteria affects the parasite’s motility and mechanical force generation, both of which are key to its pathogenicity. Using a micropillar-array system and live imaging, we found that exposure to enteropathogenic *Escherichia coli* significantly increases *E. histolytica*’s motility while reducing its traction force on fibronectin-rich matrices. These changes in behavior highlight the role of bacterial interactions in enhancing the parasite’s virulence. Our findings provide important insights into the mechanistic basis of *E. histolytica*’s pathogenicity, offering potential avenues for new treatments against amebiasis.

## Introduction

Amebiasis, caused by the protozoan parasite *Entamoeba histolytica*, is a parasitic infection that affects approximately 50 million people and causes 100,000 deaths each year worldwide, especially in regions with inadequate sanitation and hygiene practices [1, 2]. The transmission of this parasitic infection occurs via the fecal-oral route, predominantly through contaminated food or water containing *E. histolytica* cysts (the infective form). Upon entry into the host’s intestine, the cyst undergoes excystation, releasing trophozoites (the invasive form) that can invade the colonic mucosa, leading to tissue destruction and abscess formation [3, 4].

Invasive intestinal disease caused by *E. histolytica* can manifest as abdominal pain and bloody diarrhea, corresponding histologically with trophozoites invading and undermining the intestinal surface laterally, forming the characteristic flask-shaped ulcers. Rarely, *E. histolytica* trophozoites can enter the bloodstream and disseminate to other organs, most often to the liver, where they result in tissue destruction with inflammation, forming amebic liver abscesses [4, 5].

The invasiveness and virulence are largely dependent on motility and physical force exerted by *E. histolytica*, particularly through mechanisms of adherence, tissue invasion, and cytotoxic activities. The motility of *E. histolytica* is powered by its dynamic actomyosin cytoskeleton, which includes fibrous actin (F-actin) and myosin proteins [6, 7]. The parasite’s movement, particularly characterized by amoeboid migration, is facilitated not only by the formation of pseudopods at the leading edge (driven by local actin assembly), but also by contraction at the rear, generating pressure propelled by actin and myosin II. These motility properties enable the parasite to navigate through host microenvironment and tissues, i.e., intestinal mucus and epithelia, extracellular matrix (ECM), and blood vessels [8, 9]. Following the degradation of the mucus layer and the removal of epithelial cells, *E. histolytica* migrates across the fibronectin-rich basement membrane towards the crypts of Lieberkühn, before invading the lamina propria and deeper structures [10]. During this migration and invasion, *E. histolytica* trophozoites adhere to the ECM and form adhesions to fibronectin by interacting with it via a β1-integrin-like receptor [11–13]. Traction force is generated by the actin cytoskeleton and transmitted to the ECM through these adhesions [14–16].

The human intestine harbors an estimated 10^13 to 10^14 microbes across over 1,000 identified microbial species [17–19]. The interactions among these microbes, as well as between the microbes and the host, are highly complex and frequent, playing a crucial role in intestinal physiology and pathology [19]. Notably, the crosstalks between *E. histolytica*, the host digestive and immune systems, as well as the microbiota that influence amebiasis are emerging as a significant area of research [20, 21]. For instance, multiple interactions between *E. histolytica* and bacterial biofilms can affect disease persistence and antibiotic resistance [21]. The composition of the gut microbiota varies among individuals with different clinical manifestations of *E. histolytica* infection, such as asymptomatic colonization, colitis, and liver abscess [22–24]. Normal microbiota can protect the host from amebic colitis by recruiting neutrophils to the gut via the CXCR2 pathway [24, 25]. Furthermore, the metabolite deoxycholic acid produced by *Clostridium scindens* has been shown to increase granulocyte-monocyte progenitors in the bone marrow, which in turn elevates colonic neutrophil levels during amebic infection [20, 26]. Additionally, segmented filamentous bacteria, another commensal, interact with bone marrow dendritic cells through serum amyloid A, leading to the recruitment of colonic neutrophils, dendritic cells, and upregulation of interleukin-17A as part of the immune defense against amebiasis [20, 27].

However, research on the direct interactions between *E. histolytica* and other microbes is limited, highlighting the urgent need for further investigation. These interactions, accompanied by *E. histolytica* invading and destroying the intestinal mucosa layers, significantly influence the parasite’s invasiveness and virulence [19]. Several in vitro studies have demonstrated that the presence of enteropathogenic bacteria enhances the pathogenic behavior of *E. histolytica*. For instance, interaction with virulent bacteria such as enteropathogenic *Escherichia coli*, *Shigella dysenteriae*, or *Clostridium symbiosum* can increase the parasite’s cysteine proteases activity, cytopathic effect, erythrophagocytosis, hemolytic activity, and host’s proinflammatory cytokines secretion [28–31]. However, whether the interaction between bacteria and *E. histolytica* has a crucial impact on the motility and physical force of the parasites, and thus further affects *E. histolytica*’s invasiveness, remains unknown.

In this study, we employed a state-of-the-art micropillar-array system [32] combined with live imaging—a cutting-edge tool in mechanobiology—to investigate whether and how enteropathogenic *E. coli* affect the traction force exerted by *E. histolytica* on substrates. By analyzing changes in traction force, we aimed to uncover how these alterations affect the motility and invasiveness of the parasite. Our investigation seeks to illuminate the role of physical interactions and cellular behaviors in the virulence of *E. histolytica*, offering new insights into the mechanistic basis of its pathogenicity.

## Results

### *E. histolytica* generates significantly higher traction force on fibronectin-coated micropillars compared to uncoated ones

Polydimethylsiloxane (PDMS) micropillar-array is an effective system to study cellular traction forces within a range of physiological stiffnesses, including tissues that *E. histolytica* trophozoites encounter in the gut due to the excellent biocompatibility and high structural flexibility of PDMS [32–34]. The elastic PDMS micropillars allow cells to adhere to and to be bent. Tracking cell movements and pillar displacements with live-cell imaging enables the analysis of cell motility and cellular forces (Fig 1A, see Methods for details). We coated the surface micropillar-array with human fibronectin, as interaction with the ECM, particularly fibronectin, plays a paramount role in *E. histolytica* invasion and virulence [35, 36]. Indeed, histology slides from the Human Protein Atlas (HPA, www.proteinatlas.org) (37) revealed that fibronectin is abundant in human intestinal mucosa, with particularly prominent expression in the subepithelial basement membrane and lamina propria (S1 Fig).

**Fig 1.**
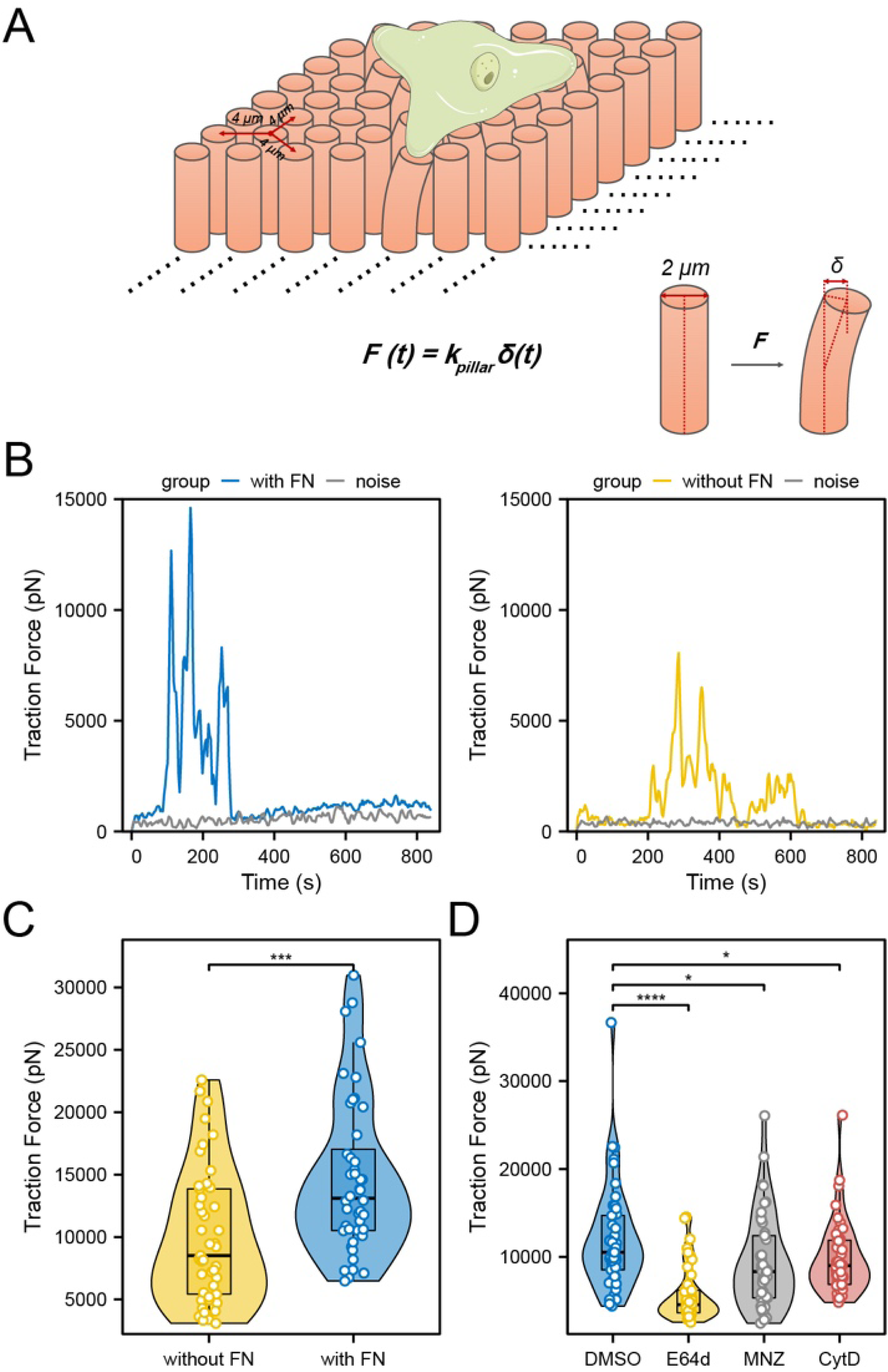
*E. histolytica* generates significantly larger traction force on fibronectin-coated micropillars compared to uncoated ones. (A) Schematic diagram showing how a trophozoite interacts with the micropillar-array surface and thereby generating traction force, leading to the deflection of micropillars. The traction force as a function of *F(t)* is calculated by multiplying the spring constant of a micropillar *k_pillar_* and micropillar displacement over time *δ(t)*. The cartoon illustration of the *E. histolytica* trophozoite was adopted and modified from Servier Medical Art. (B) Representative graphs showing the temporal changing of traction force on a micropillar as a trophozoite passing on a micropillar with (left) or without (right) fibronectin (FN) coating. The grey curve indicates background noise of micropillars untouched by the cells. (C) Comparison of the max traction force on micropillars exerted by *E. histolytica* between fibronectin-coated (n = 44) and uncoated micropillars (n = 45). (D) Comparison of the max traction force on fibronectin-coated micropillars exerted by *E. histolytica* between trophozoites treated with DMSO (control) and cysteine protease inhibitor E64d, metronidazole (MNZ), or F-actin polymerization inhibitor Cytochalasin D (CytD). The analyses included 45 micropillars for each treatment condition. * *P* < 0.05, *** *P* < 0.001, **** *P* < 0.0001.

We have tested several arrays of micropillars with the same diameter, 2 μm, and different heights including 5.3, 9.4, and 13.2 μm, corresponding to rigidities (spring constant, *k_pillar_*) of 2, 6 and 31 pN/μm, respectively, calculated by Euler-Bernoulli beam theory [38]. Consequently, only micropillars-array of height 5.3 μm was employed in this study, since *E. histolytica* applies relatively strong forces that lead to too large micropillars deflections on softer/higher micropillars-arrays (9.4 or 13.2 μm height), which resulted in tracking difficulty for later analyses. The effective elastic modulus of the micropillars-array of height 5.3 μm is approximately 22.7 kPa, which is within the human physiological range [39].

To assess the biomechanical relevance of this system, we first compared the traction force applied by *E. histolytica* trophozoites on pillars with and without fibronectin coating. Curves of traction force as a function of time *F(t)* of specific micropillars were plotted showing the magnitude and characteristics of cellular force (Fig 1B). For comparisons between different conditions, we used the maximum force value from each curve, representing the highest traction force exerted by the cell on the corresponding micropillar. The results show that *E. histolytica* trophozoites apply significantly greater traction force on fibronectin-coated micropillars compared to uncoated ones. (Fig 1B and 1C).

Analyses of the motility properties of *E. histolytica* on the micropillars, including measuring the velocity, Euclidean distance, and directionality of *E. histolytica*, showed no significant differences between coated and uncoated micropillars (S2 Fig).

Next, we employed three inhibitors and/or medications for *E. histolytica* that are expected to affect cellular force generation on fibronectin-coated micropillars by influencing cell adhesion and the actin cytoskeleton. The treatment concentrations for *E. histolytica* were determined based on published literature [40–42]. Only visibly active migrating cells observed in the live-imaging videos were included in the analysis. We applied the F-actin polymerization inhibitor Cytochalasin D (CytD) since traction force generation depends on F-actin cytoskeleton contraction and adhesion formation on ECM [14–16]. We used the cysteine protease inhibitor E64d since E64 derivatives can significantly decrease the adhesion of *E. histolytica* to ECM proteins [43]. We also applied the antibiotic metronidazole (MNZ), a drug currently used to treat amebiasis, as its treatment induces the formation of oxidized proteins in *E. histolytica*, including cytoskeletal proteins such as actin. This oxidative stress not only impairs the formation of F-actin [42], but also reduces the adhesion of *E. histolytica* to substrate [44].

Traction forces exerted by *E. histolytica* on fibronectin-coated micropillars significantly decreased when treated with these inhibitors (Fig 1D). Taken together, depriving the ECM component fibronectin from substrate, preventing F-actin polymerization (CytD), suppressing cysteine protease (E64d), and inducing oxidative stress (MNZ), all significantly reduce traction force of *E. histolytica* on the external substrate.

### Exposure to enteropathogenic *E. coli* led to significant alterations in both motility properties and cellular contraction force of *E. histolytica*

*E. histolytica* resides in a bacteria-rich environment within the intestine, where it not only interacts with but also feeds on various bacterial species, which in turn can modulate its virulence [45, 46]. The cytopathic assay confirmed that a short period (30 minutes) of co-incubation of trophozoites with *E. coli* resulted in a greater percentage of destruction of a HeLa cell monolayer, with the enteropathogenic O55 strain showing the highest ability in enhancing the parasite’s destructive effect (S3 Fig).

To better replicate the conditions *E. histolytica* trophozoites encounter during the invasion along fibronectin-rich basement membrane beneath the human intestine epithelia, as well as the further invasion to the lamina propria (S1 Fig), our subsequent experiments were all conducted using human fibronectin-coated micropillars.

To investigate the impact of bacterial exposure on the invasiveness of *E. histolytica*, which is partly manifested through changes in motility, we pre-incubated the trophozoites with *E. coli* (enteropathogenic O55 or nonpathogenic K12) at 37°C for 30 minutes, using the same conditions as the previously described cytopathic assay. This was followed by live imaging using the micropillar system. We then analyzed and compared the motility features of trophozoites under three conditions: 1) control (no bacterial exposure), 2) incubation with *E. coli* O55, and 3) incubation with *E. coli* K12. Analysis of movement trajectories revealed that trophozoites exposed to bacteria migrated significantly farther than the control group, with those incubated with *E. coli* O55 showing the greatest distance traveled (Fig 2A; S1 and S2 Video). Further quantitative analyses indicated that the directionality and Euclidean distance of trophozoites incubated with *E. coli* O55 were significantly higher than control (Fig 2B and 2C). The motility characteristics of trophozoites incubated with *E. coli* K12 were intermediate between the control and those incubated with *E. coli* O55. The velocity distribution of trophozoites incubated with *E. coli* O55 appears higher than that of control group, but did not reach statistical significance (*P* = 0.141) (Fig 2D).

**Fig 2.**
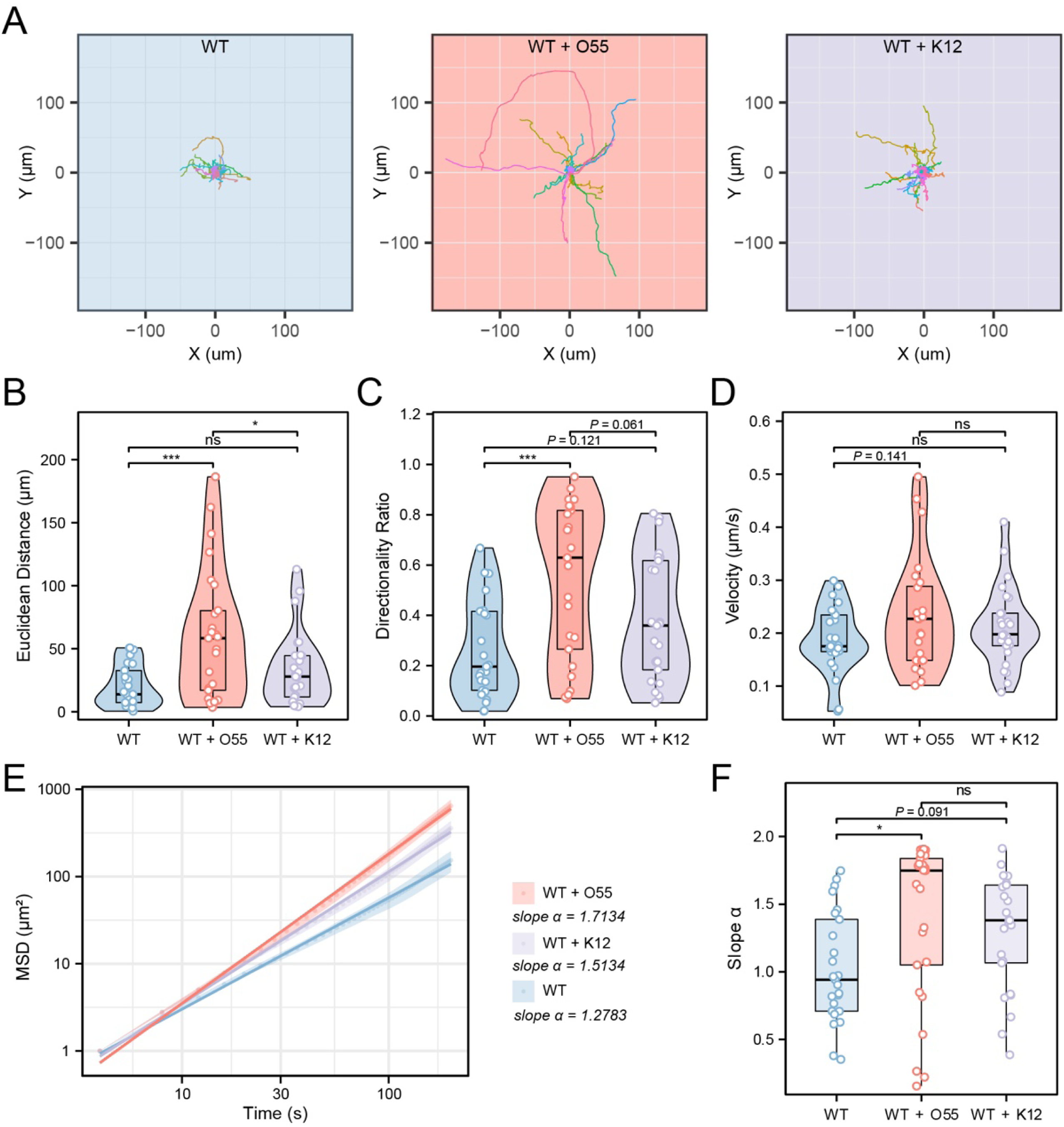
Significant differences in motility properties of *E. histolytica* following short period of *E. coli* exposure. (A) Trajectory plots regarding trophozoite migration under three conditions: control, incubation with enteropathogenic *E. coli* O55, and incubation with nonpathogenic *E. coli* K12. (B-D) Quantitative comparison of migration Euclidean distance (B), directionality (C), and velocity (D) of *E. histolytica* under the different conditions described in (A). (E) The relationship between mean square displacements (MSD) and the corresponding time data was averaged and plotted on a logarithmic scale for *E. histolytica* locomotion under the different conditions. (F) Comparison of the slopes (α) for each individual cell/plot under the different conditions. For all the above diagrams, n = 25 in each group. Notably, the average slope α of O55 group is the highest, indicating that it exhibits the strongest Lévy-like movement pattern. α = 2: ballistic motion, α > 1: superdiffusion (Lévy-like movement), α = 1 for diffusive/Brownian motion. * *P* < 0.05, *** *P* < 0.001.

To better understand the cell migration mode, we calculated the mean square displacements (MSD) of the cells, a well-established method in cell migration analysis [47, 48], including in *E. histolytica* studies [49, 50]. MSD measures the surface area explored by cells over time, reflecting migration efficiency. It provides insights into both speed and directional persistence and is typically shown in a log(MSD)-log(time) plot [47]. The slope of this plot, α, indicates the motion type: α = 2 for ballistic motion (persistent in direction), α > 1 for superdiffusion (Lévy-like movement), α = 1 for diffusive/Brownian motion (random movement), and α < 1 for subdiffusion (restricted movement) [47–49, 51].

Analyses of the averaged log(MSD)-log(time) plots across all cells (Fig 2E), along with the slope α-values for individual cells (Fig 2F), demonstrated that trophozoites incubated with *E. coli* O55 exhibited the highest α-value (α = 1.7134), followed by those incubated with *E. coli* K12 (α = 1.5134) and the control group (α = 1.2783). This suggests that in the control group, *E. histolytica* trophozoites exhibit movement close to diffusive or Brownian motion, whereas those incubated with *E. coli* O55 display superdiffusion or a Lévy movement mode, characterized by significantly greater directionality and velocity.

To explore whether cellular traction force can be affected following *E. coli* exposure, we further analyzed and compared the traction force on micropillars by *E. histolytica* under the three conditions mentioned above: control, trophozoites incubated with enteropathogenic *E. coli* O55, and trophozoites incubated with nonpathogenic *E. coli* K12. This revealed that trophozoites incubated with *E. coli* O55 exhibited the lowest traction force, followed by those incubated with *E. coli* K12, whereas the control group displayed the highest traction force on the micropillar- arrays (Fig 3A and 3B).

**Fig 3.**
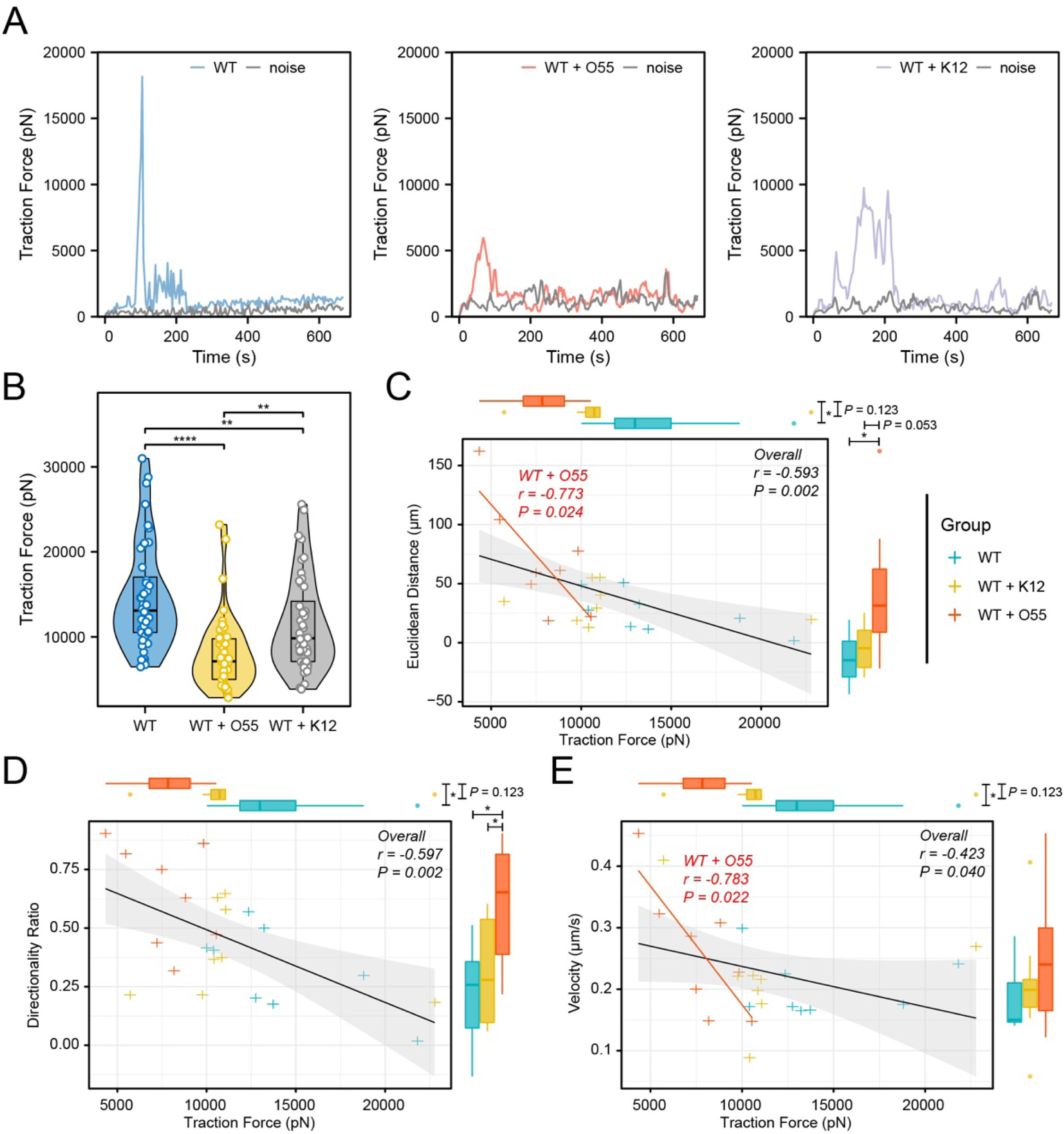
Significant difference of cellular traction force on micropillars of *E. histolytica* following short period of *E. coli* exposure. (A) Representative graphs showing the changing in traction force exerted on micropillars over time for control trophozoites (left), trophozoites incubated with enteropathogenic *E. coli* O55 (middle), and trophozoites incubated with nonpathogenic *E. coli* K12 (right), during a trophozoite passing on a micropillar. The grey curve indicates background noise of micropillars untouched by the cells. (B) Quantitative comparison of the max traction force on micropillars exerted by *E. histolytica* between the different conditions: control (n = 44), *E. coli* O55-incubated (n = 43), and *E. coli* K12-incubated (n = 44) trophozoites. (C-E) Correlation between cell motility indicators and traction force; n = 8 in each group. * *P* < 0.05, ** *P* < 0.01, **** *P* < 0.0001.

The trend in cellular traction force and that in motility characteristics across the three *E. histolytica*-*E. coli* co-incubation conditions were opposite, prompting further consideration of the relationship between cellular traction force and motility. In the group without exposure to *E. coli*, the cells exhibited a non-directional exploratory behavior, actively using pseudopods to tug on the surrounding pillars. At the same time, the cells seemed to exhibit greater adhesion, resulting in more significant deflection of the micropillars (S3 Video). However, when exposed to *E. coli*, the cells transitioned to an enhanced Lévy-like movement pattern (rapid and goal-directed), randomly exploring and displaying noticeably smaller deflections of the micropillars (S4 Video). To illustrate the exact correlation between cell motility and traction force quantitively, linear regressions and Pearson correlations were performed for cells from *E. histolytica*-*E. coli* co-incubation conditions. This showed that all three cell motility indicators are negatively correlated with traction force, with Euclidean distance and directionality ratio exhibiting a high negative correlation (r < -0.5) (Figs 3C-E). The data of trophozoites incubated with *E. coli* O55 showed a higher negative correlation with r < -0.7 (Fig 3C and 3E).

### Exposure to enteropathogenic *E. coli* altered the F-actin morphology and spatial localization in *E. histolytica*

Given that co-incubation of enteropathogenic *E. coli* led to remarkable variations in the motility and cellular traction force of *E. histolytica*, we next sought to investigate whether the actin cytoskeleton plays a crucial role. To explore how *E. coli* exposure affects F-actin organization in *E. histolytica* control trophozoite, trophozoites incubated with *E. coli* O55, and trophozoite incubated with *E. coli* K12 were seeded on fibronectin-coated micropillars and incubated at 37°C for one hour, followed by fixation and subsequent staining with AF-488 Phalloidin. Notably, all images scanned by confocal microscopy were focused on the plane of the micropillars’ top surface. Consequently, a significant reduction of F-actin intensity in *E. histolytica* following *E. coli* exposure was observed, with no significant difference found between trophozoites incubated with *E. coli* O55 and *E. coli* K12 (Fig 4A and 4C).

**Fig 4.**
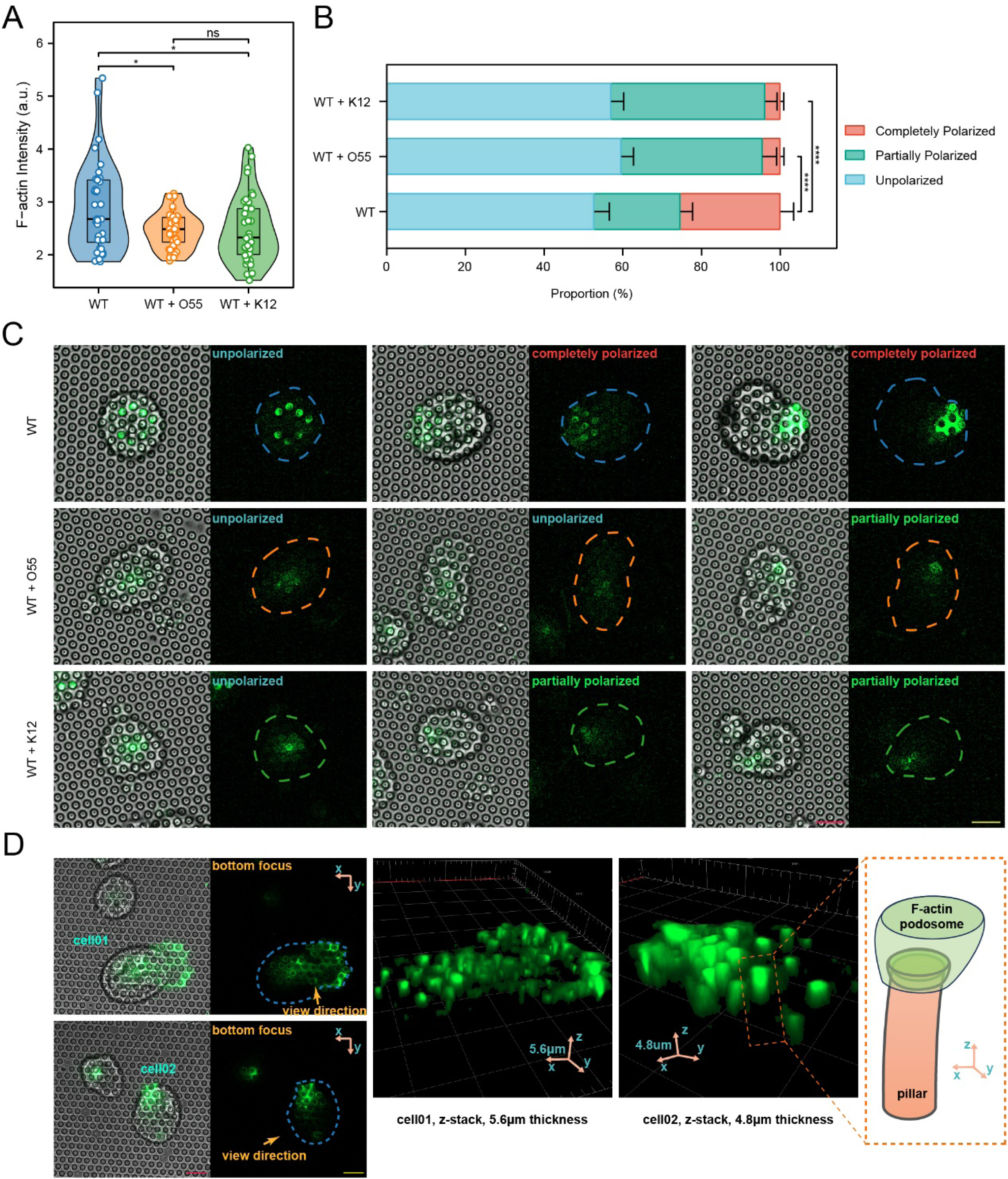
Alteration of F-actin morphology in *E. histolytica* following *E. coli* short-period exposure. (A) Quantitative comparison of Phalloidin intensity in control (n = 40), enteropathogenic *E. coli* O55-incubated (n = 41), and *E. coli* K12-incubated (n = 40) *E. histolytica* trophozoites. \**P* < 0.05. (B) Comparison of the proportion of F-actin- polarized cells in control (n = 160), *E. coli* O55-incubated (n = 202), and *E. coli* K12- incubated (n = 209) *E. histolytica*. (C) Representative confocal images (20x objective) showing the Phalloidin signal in *E. histolytica* on micropillars, both with polarized and unpolarized samples. (D) Left panel: high-resolution confocal images (63x objective) captured by focusing on the plane of micropillar-array top surface, showing that F-actin formed rings that tightly encircle the corresponding micropillars, as well as filaments linking the rings. Right panel: z-stack reconstruction of F-actin spatial distribution near the micropillars, presenting podosome-like structure. The schematic diagram in the dashed-line frame illustrates the spatial relationship between an F-actin podosome-like structure and its corresponding micropillar. Scale bar in C and D: 10 μm. * *P* < 0.05, **** *P* < 0.0001.

To provide additional evidence, we performed Phalloidin staining on cells plated in ibidi µ-Slide 8 Well plates. F-actin staining of *E. histolytica* trophozoites on fibronectin-coated substrates showed significantly higher intensity compared to those on uncoated micropillars (S4 Fig A and E). Moreover, on the fibronectin-coated substrate, *E. histolytica* trophozoites exposed to *E. coli* O55 exhibited significantly lower F-actin intensity than those exposed to K12 and the control group (S4 Fig C and E), consistent with the results observed on the micropillars. These findings align with previous results, showing that *E. histolytica* trophozoites exerted greater traction forces on fibronectin-coated micropillars compared to uncoated ones (Fig 1B and 1C), as well as decreased traction forces when exposed to *E. coli* (Fig 3A and 3B). The positive relation between F-actin intensity at micropillar top surface and cellular traction force suggests that elevated traction force on micropillars may be dependent on enhanced F-actin polymerization at pseudopods.

Given that the intracellular local polarization of F-actin is closely correlated with traction force on micropillars [38], we further analyzed the polarity of the Phalloidin signal in different conditions, categorizing it into completely polarized (towards the cell edge), partially polarized (off-center but not reaching the cell edge), and unpolarized (centered) F-actin distributions (Fig 4B and 4C). This analysis showed a significant reduction in the proportion of completely polarized cells when exposed to *E. coli* O55 and K12 (Fig 4B and 4C). This F-actin polarization pattern is consistent with the Phalloidin intensity comparisons between trophozoites incubated with *E. coli*.

Finally, we analyzed the spatial relationship between F-actin and its corresponding micropillars using high-resolution confocal scanning combined with the z-stack reconstruction mode. We discovered that F-actin accumulated near the micropillars, forming podosome-like adhesion structures that extend downward, wrapping around the top portion of the micropillars (Fig 4D, right panel). When observed from the plane of the micropillar’s top surface, F-actin formed rings that tightly encircle the corresponding micropillars, as well as filaments linking the rings (Fig 4D, left panel). This arrangement helps concentrate the force for gripping a micropillar, enabling more effective application of traction force to the micropillar [38]. Next, we examined the spatial localization difference of F-actin between three conditions: *E. histolytica* control trophozoite, trophozoites incubated with *E. coli* O55, and trophozoite incubated with *E. coli* K12, which might shed light on the mechanism regarding the negative correlation between cell locomotion and traction force on micropillars. It can be observed that without exposure to *E. coli*, F-actin primarily accumulated at podosomes (Fig 5A and 5B), possibly enabling the cells to grip the micropillars firmly and effectively. Additionally, F-actin tended to polarize toward pseudopods at the cell edge. However, upon exposure to *E. coli*, a significant amount of F-actin shifted to the upper part of the cytoplasm (Fig 5A and 5C). Consequently, the F-actin signal at the podosomes corresponding to the micropillars weakened, resulting in shorter and less prominent adhesions. There was a significant negative Pearson correlation (r = -0.458, *P* < 0.0001) between the height of F-actin structures in the upper part of cell and the height of adhesion structures (Fig 5D).

**Fig 5.**
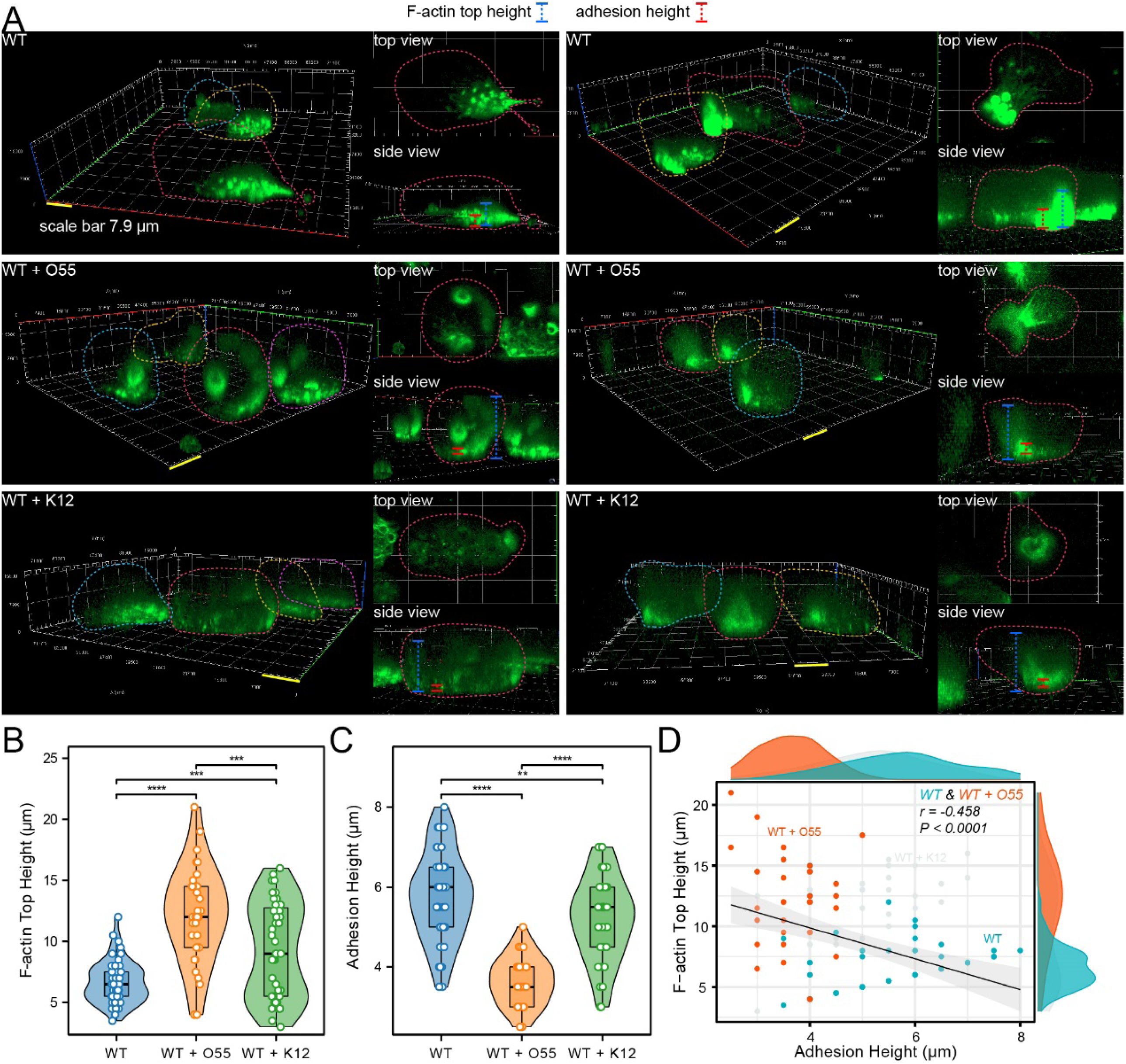
The spatial localization difference of F-actin between following *E. coli* short-period exposure. (A) Z-stack representative images showing the spatial localization differences of F-actin among three conditions: *E. histolytica* control trophozoites (n = 54), trophozoites incubated with enteropathogenic *E. coli* O55 (n = 32), and trophozoites incubated with *E. coli* K12 (n = 43). (B) Comparison of the height of F-actin structures in the upper part of the cell, distinct from adhesion structures on micropillars, under the different conditions. (C) Comparison of the height of adhesion structures under the different conditions. (D) Pearson correlation between the height of F-actin structures (in the upper part of the cell plasma) and the height of adhesion structures.

Additional analysis based on Phalloidin staining of *E. histolytica* trophozoites on fibronectin-coated ibidi µ-Slide 8 Well plates revealed that when trophozoites were incubated with *E. coli* O55, the ratio of basal surface area (determined by confocal F-actin imaging) to the total area in the bright field (non-confocal imaging) was significantly smaller compared to control trophozoites. This suggests that interaction with *E. coli* O55 reduces the contact surface area with the fibronectin-coated substrate (S4 Fig D and E).

## Discussion

*E. histolytica*, a parasite that coexists with and feeds on bacteria in the intestinal environment, demonstrates motility closely linked to its invasiveness [6]. Despite the known short-period co-incubation of *E. coli* (O55 and K12) and *E. histolytica* trophozoites can promote *E. histolytica* cytotoxicity [45, 52, 53], the impact of bacterial presence on *E. histolytica*’s motility and physical forces, and its subsequent effect on invasiveness, remains underexplored.

In this study, we employed a micropillar-array system with live imaging, a technique designed for investigating traction forces on the substrate due to cellular contractility in mechanobiology [32], to explore the effect of *E. coli* (especially enteropathogenic *E. coli*) exposure on *E. histolytica* migration. Our approach involved assessing the traction force, motility characteristics (including velocity, Euclidean distance, directionality ratio, and migration mode), and F-actin spatial morphology of *E. histolytica* under conditions with and without two strains of *E. coli*. Our results indicate that *E. histolytica* modifies its physical properties and motility in response to bacterial presence, demonstrating an interplay between bacterial presence and the dynamics of the parasite’s actin-rich cytoskeleton. These findings contribute to a better understanding of the role of bacteria in modulating cellular behavior and pathogenicity.

*E. histolytica* invasion of the intestine begins with the removal of the protective mucus layer of the epithelium, facilitated by the parasite’s cysteine proteases, such as CP-A5. Following this, the trophozoites adhere to and detach enterocytes, leading to the disruption and degradation of the epithelial barrier [2]. Interestingly, the pathogenic trophozoites do not indiscriminately invade the lamina propria. Instead, they prefer to migrate along the fibronectin-rich basement membrane deep into the crypts of Lieberkühn (first stage), where they eventually penetrate the lamina propria (second stage) [2, 10, 54]. Using histology slides from the HPA, we highlighted that both the basement membrane and lamina propria of the human large intestine are enriched with fibronectin. Therefore, in the present study, we used a fibronectin-coated substrate (micropillar arrays and Ibidi plates) to mimic these pathological scenarios. In the first stage, *E. histolytica* invades parallel to the fibronectin-rich basement membrane rather than invading perpendicular to it. Thus, the invasiveness of *E. histolytica* is primarily characterized by its directional invasion along the basement membrane, allowing it to reach deep inside the crypts of Lieberkühn more rapidly. In the second stage, *E. histolytica* seeks to invade a broader area as quickly as possible, ultimately leading to the formation of a flask-shaped ulcer [1]. This virulent invasion is partly mimicked in *E. histolytica*’s cytopathic activity to cause extensive destruction of the HeLa cell monolayer in a period of time. Taken together, the invasive behavior highlighted here is the rapid and directional movement of *E. histolytica* along the fibronectin-coated plane, resulting in efficient invasion to a greater distance and a broader area.

Unlike mesenchymal motility, amoeboid migration is characterized by the formation of blebs and pseudopods at the front of the trophozoite. This process, driven by hydrostatic pressure originating from the rear, involves a retraction- relaxation cycle in which the actomyosin cytoskeleton and actin-binding proteins retract the cell body and push the trophozoite forward [54–56]. Besides the role in bleb formation, F-actin is also responsible for developing adhesion structures including podosome-like, stress fiber-like structures, and adhesion plates when *E. histolytica* is placed on fibronectin substrates and interacts with fibronectin via a β1- integrin-like receptor [11–13, 54].

We assume that both amoeboid migration and adhesion to fibronectin substrate contribute to the displacement of micropillars. Without exposure to *E. coli*, *E. histolytica* trophozoites exhibited non-directional exploratory behavior (localized wandering/rotating), actively using pseudopods to pull on the surrounding pillars. Plus, the cells seemed to exhibit greater adhesion. This locomotion style results in more significant deflection of the micropillars. However, when exposed to enteropathogenic *E. coli*, the cells transitioned to an enhanced Lévy-like movement pattern (rapid and directional), without random exploration and active micropillar pulling, leading to a noticeably smaller deflection of the micropillars.

Given that cellular traction force on micropillars is found negatively correlated with the motility (directionality and velocity) of *E. histolytica* in the present study, it can be speculated that once interacting with enteropathogenic *E. coli*, the trophozoites focus more on rapid directional movement, thereby reducing the traction force and adhesion strength exerted on the underlying micropillars. This may be related to changes in the spatial dynamics and localization of F-actin structures.

Supporting this, we found that without exposure to *E. coli*, F-actin primarily accumulated at podosomes, which were notably large and elevated, possibly enabling the cells to grip the micropillars firmly and effectively. Additionally, F-actin tended to polarize toward pseudopods at the cell edge. However, upon exposure to enteropathogenic *E. coli*, a significant amount of F-actin shifted to the upper part of the cytoplasm in many cells. Consequently, the F-actin signal at the podosomes corresponding to the micropillars weakened, resulting in shorter and less prominent adhesions. There was a significant negative correlation found between the height of F-actin structures in the upper part of the cells and the height of adhesion structures. We and others have found that F-actin is highly concentrated not only at the adhesion structures on the bottom and actomyosin cytoskeleton cortex at periphery, but also in the middle of the cell [55]. Upon exposure to *E. coli*, it is possible that the shift of F-actin towards the upper part of the cytoplasm is linked to phagocytosis, potentially contributing to phagosome formation. Cell motility and phagocytosis are widely recognized as inseparable processes [6, 57], sharing common features in mechanics and morphology, as well as key elements in biological process and cellular component, including actin, Myosin IB, Myosin II, and PAK [6, 57–61]. Indeed, motility and phagocytosis are likely intertwined processes that evolved in tandem, reflecting an inseparable function. The ability to actively move towards bacterial prey would have provided a significant evolutionary advantage over passively waiting for food to arrive, enhancing competition for ingestion [57].

Hence, the F-actin structure shifted to the upper part of the cytoplasm during *E. coli* exposure is likely involved in the rapid and direction-persistent amoeboid migration of *E. histolytica*, which is not necessarily driven by cellular active traction force on the substrate but primarily by hydrostatic pressure generated by the rapid contraction of actomyosin machinery at the rear [54, 62, 63]. In contrast, without *E. coli* exposure, the F-actin podosome near the micropillar surface is more dominant and primarily responsible for deflecting the pillars, and, together with the less dominant hydrostatic-powered blebs, contributes to the exploratory motility style.

There are several explanations to the negative correlation between traction force on micropillars and the motility (directionality and velocity) of *E. histolytica*. First, the enhanced Lévy-like movement requires a large accumulation of F-actin at the rear, which competes with the F-actin used in the construction of podosomes. Remarkably, the motility enhancement is powered by enteropathogenic *E. coli* exposure, which may provide targets for cells to pursuit. Phagocytosis might contribute to the shift of F-actin towards the upper part of the cytoplasm, as simply reducing cell adhesion without fibronectin coating does not significantly alter motility behavior. Second, the rapid and directional movement results in less time spent on a single pillar, making it difficult to form stable podosomes, thus reducing cell’s adhesion to micropillars. Third, from a kinematic perspective: the driving force (*F*) of forward movement arises from the force and counterforce generated by the rapid and directional motion, while the resistance (*f*) stems from the adhesion between the cell and micropillars. When maximum velocity and power are reached, *P_max_ = F · v_max_*, while *F = f*. Hence, to actively increase invasion efficiency and enhance *v_max_*, *f* needs to be decreased, leading to reducing micropillar deformation, which is achieved by weakening adhesion, as well as reducing the contact area on the substrate.

A newly published study by Manich et al. [54] explored the impact of fibronectin on *E. histolytica* trophozoite motility, with a focus on fibronectin-induced changes in morphodynamics and an increase in adhesion force. This is consistent with our findings that fibronectin-coating enhances cell’s traction force on micropillars, which is dependent on a corresponding strengthen of cell adhesion, as evidenced by enhanced F-actin intensity in the bottom cell plane under fibronectin condition (micropillar array or ibidi plate). Manich et al. also reported significantly faster migration on glass compared to fibronectin-coated surfaces, which is consistent with our finding of a negative correlation between cell motility and substrate adhesion. However, the resistance to cell forward movement might differ between micropillar arrays and flat plates (friction), which is also reflected by the adhesion morphology. It is also worth mentioning the difference in stiffnesses between glass plate, which is extremely rigid, and the micropillar-array system, which mimics physiological stiffnesses. Based on the above findings, we further demonstrate that exposure to enteropathogenic *E. coli* makes *E. histolytica* reduce the adhesion and traction force on micropillars, while enhancing the Lévy-like movement.

It is worth noting that the comparison of velocity between *E. histolytica* incubated with O55 and the control group did not reach statistical significance at the 0.05 level. However, we observed differences in the data distribution for velocity. Velocity calculation based on manual tracking of cells is limited by the difficulty of accurately determining the centroid due to their dynamic and changing morphology. This may have led to an underestimation of the velocity differences, introducing potential measurement bias. Additionally, two other analyses support the finding that O55 co-incubation increases cell velocity: 1) Traction force and velocity are significantly negatively correlated, particularly in the O55 co-incubation group, where the correlation coefficient (r) reaches -0.78, the strongest correlation observed between motility parameters and traction force. 2) The migration mode analysis, which utilizes α values from the log(MSD)-log(time) plot to account for both velocity and directionality [47], revealed statistically significantly higher α values (enhanced Lévy-like movement) in the O55 co-incubation group.

It is of great value to further investigate the mechanism regarding what interaction occurs between *E. histolytica* and enteropathogenic *E. coli*, driving a motility enhancement. We speculate that this may be due to the *E. histolytica* chasing and engulfing bacteria, which expands the *E. histolytica*’s invasion range. Kinematic studies on cancer metastasis demonstrate that the trajectories of metastatic cells exhibit also enhanced Lévy-like movement patterns, characterized by clusters of small steps interspersed with long ’flights’. In contrast, non-metastatic cancer cells move diffusively. Lévy-like movement is also employed by animal predators searching for sparse prey. This migration mode might provide metastatic cells with an effective strategy for disseminating and searching suitable sites to establish new metastatic sites, which is a similar goal for *E. histolytica* [48].

We co-incubated *E. histolytica* with two strains of *E. coli*, but the outcomes differed between the two strains. O55 is classified as an enteropathogenic *E. coli* [64, 65]. However, K12 is not enteropathogenic, but a laboratory strain that has undergone extensive mutagenesis, leading to the loss of several genetic traits associated with virulence [66, 67]. *E. histolytica* co-incubated with O55 not only significantly increased *E. histolytic*’s destruction of cultured HeLa cell monolayers, but also markedly enhanced Lévy-like movement, while significantly reducing its traction force on the fibronectin-rich substrate. In contrast, the cytopathic activity of *E. histolytica* and the associated changes in motility and cellular mechanics were less pronounced when co-incubated with K12 (compared to O55). These findings highlight a clear link between the parasite’s motility/traction force and cytopathic activities, suggesting that changes in kinematics and mechanics play a pivotal role in the virulence and pathogenicity of

*E. histolytica*. It is reported that the galactose and N-acetylgalactosamine (Gal/GalNAc) residues on the surface lipopolysaccharide of O55 can be recognized by and interact with *E. histolytica*’s Gal/GalNAc lectin [68, 69]. In contrast, the interaction between K12 and *E. histolytica* may rely on different mechanisms, such as type 1 fimbriae mediated binding to mannose-containing receptors on the *E. histolytica* surface [68, 70]. These distinct molecular interactions may influence F-actin cytoskeleton-associated cellular components and pathways, thus affecting the locomotion, mechanics, and phagocytosis of *E. histolytica*.

In conclusion, short-term co-incubation with enteropathogenic *E. coli* enhances *E. histolytica*’s Lévy-like movement, characterized by increased directionality and velocity. This activated motility is accompanied by a reduction in F-actin-dependent traction forces and podosome-like structures on the fibronectin-rich substrate, but with increased F-actin localization in the upper part of the cytoplasm. This locomotion leads to rapid and broader dissemination, which can promote the invasiveness of *E. histolytica* (Fig 6). This study provides new insights into the role of physical behaviors in the virulence of *E. histolytica* and offers new perspectives for the clinical prevention and treatment of amebiasis.

**Fig 6.**
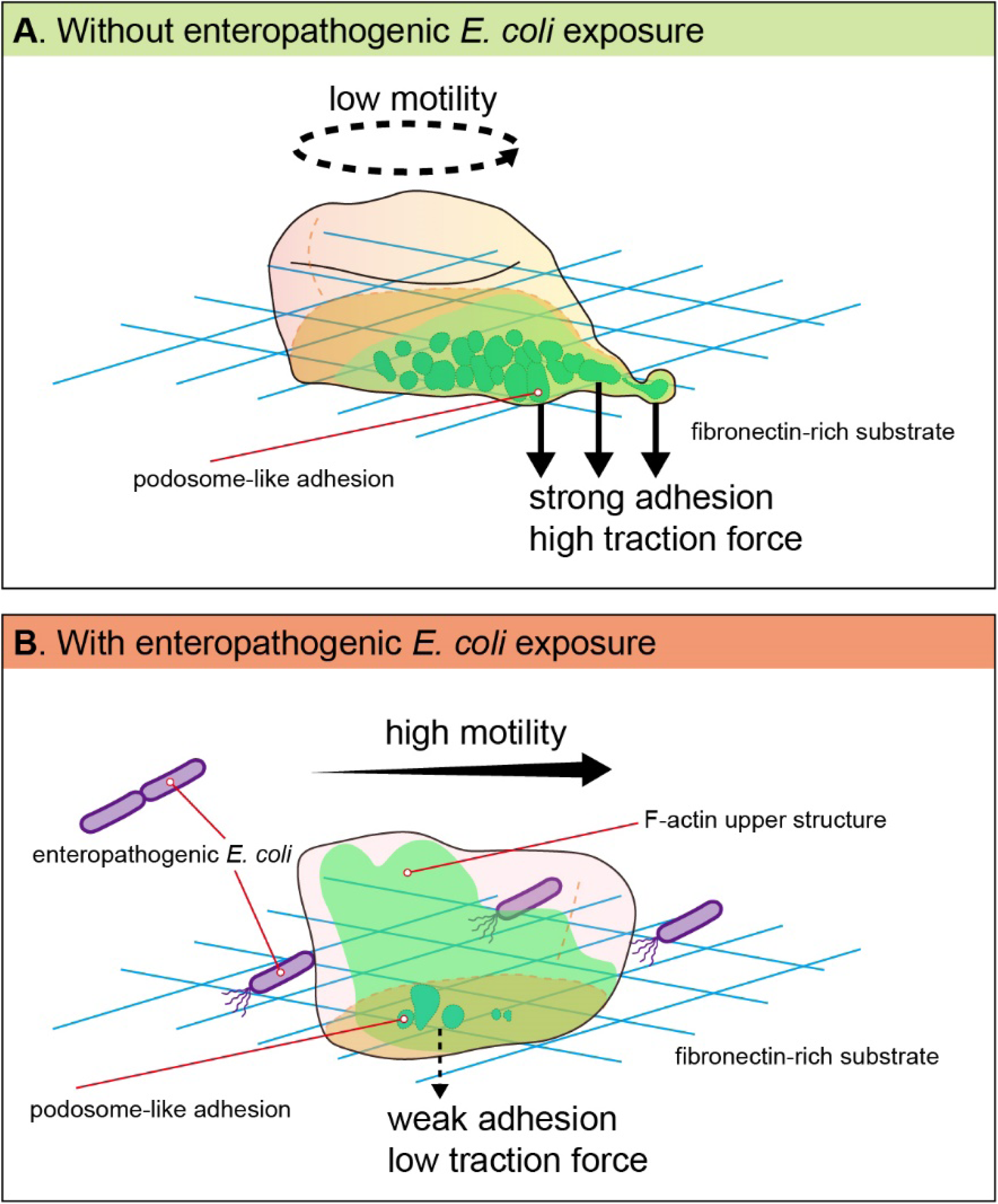
A hypothetical model illustrating the alteration in motility and traction force on a fibronectin-rich substrate during *E. histolytica* trophozoite migration with and without exposure to enteropathogenic *E. coli*. (A) Without enteropathogenic *E. coli* exposure, the *E. histolytica* trophozoites exhibit a non-directional exploratory behavior, actively using pseudopods to tug on the surrounding fibronectin-rich substrate. F-actin primarily accumulates at podosome-like adhesion structures extending downward, which are notably large, elevated and strong, possibly enabling the cells to grip the substrate firmly and effectively. Additionally, F-actin polarizes toward pseudopods at the cell edge, exerting high traction force to the fibronectin-rich substrate. (B) Co-incubation with enteropathogenic *E. coli* enhances *E. histolytica* trophozoites’ motility, characterized by enhanced Lévy-like movement patterns (increased directionality and velocity). This increased motility is accompanied by a reduction or weakening of F-actin podosome-like adhesion structures and their cell-edge polarization, as well as a decrease in traction forces on the fibronectin-rich substrate. At the same time, there is increased F-actin localization in the upper part of the cytoplasm, which may be linked to the active motility of the trophozoites. This locomotion leads to rapid and broader dissemination, which can promote the invasiveness of *E. histolytica*.

## Materials and methods

### E. histolytica strains

The *E. histolytica* strain HM-1:IMSS was graciously supplied by Prof. Samudrala Gourinath from Jawaharlal Nehru University, New Delhi, India. Trophozoites of *E. histolytica* strain HM-1:IMSS was cultured at 37°C in 13 × 100 mm screw-capped Pyrex glass tubes containing Diamond’s TYI-S-33 medium until they reached the exponential growth phase. Trophozoites were then harvested from the culture tubes using the previously outlined method of tapping followed by centrifugation, as detailed in prior protocols [71].

### Bacterial strains

The enteropathogenic *E. coli* strain TW04062 O55 is from the Thomas S. Whittam STEC Center. The nonpathogenic *E. coli* strain MG1655 K12 was provided by Sima Yaron from the Department of Biotechnology and Food Engineering at the Technion. *E. coli* cultures were incubated at 37°C in Luria-Bertani (LB) medium.

### Measurement of cytopathic activity

The assessment of trophozoites’ destruction of cultured HeLa cell monolayers was conducted utilizing a previously outlined methodology [45]. In brief, a concentration of 1 × 10^6 trophozoites/mL was incubated with *E. coli* O55 or *E. coli* K12 (1 × 10^9/mL) in serum-free Diamond’s TYI-S-33 medium at 37°C for 30 minutes while shaking. Afterwards, most bacteria were removed by washing the trophozoites with TYI medium through centrifugation at 726 × g for three cycles. Then trophozoites (10^5 per well) were cultured in serum-free Diamond’s TYI-S-33 medium and then incubated with HeLa cell monolayers in 24-well tissue culture plates at 37°C for 60 minutes. The incubation process was stopped by transferring the plates to ice, followed by removal of unattached trophozoites through washing with cold phosphate-buffered saline (PBS). The remaining HeLa cells adhered to the plates were subsequently stained with methylene blue solution (0.1% in 0.1 M borate buffer, pH 8.7). Extraction of the dye from stained cells was performed using 0.1 M HCl, and the color intensity of the extracted dye was measured spectrophotometrically at OD660.

### Micropillar-array system and traction force calculation

PDMS micropillar-arrays were employed in this study, which allowed us to detect and measure the traction force that cells applied on the micropillars [32, 38]. Based on the microscope time-series imaging of micropillars within a particular period, we spatiotemporally tracked and recorded the location of every micropillar, which was then calculated to micropillar displacement (*δ*) and then transferred to traction force (*F*) [32] (Fig 1A).

The rigidity of PDMS micropillar-array is only altered by changing micropillar height, while the cross-sectional area and chemical properties of the micropillars remain constant. The diameter of the circular, micropillars cross-section is two μm, and the center-to-center spacing between adjacent micropillars is four μm.

Tracking of micropillars movements over time was performed with NanoTracking plugin of Fiji/ImageJ (National Institutes of Health), the detailed algorithm of which is described previously [72]. Briefly, it employed a cross-correlation technique that enabled the software to obtain the relative x and y position of every micropillars in every frame of the time-series movie in nanometer-level. We only selected for analysis the cell migration-associated micropillars that were not in contact with any cells (zero position) at the beginning of the movie. Micropillar displacement (*δ*), which is caused by traction force (*F*) when cell edge/body contact the micropillars, at each frame, is calculated based on the distance from current to the zero position. Traction force (*F*) is calculated by multiplying micropillar displacement (*δ*) by the spring constant of the corresponding pillar height (Fig 1A). Micropillar arrays with heights of 5.3, 9.4, and 13.2 μm have external rigidities (spring constant, *k_pillar_*) of 2, 6, and 31 pN/μm, respectively, as calculated using the Euler-Bernoulli beam theory [38]. The curve of traction force as a function of time *F(t)* of a micropillar was plotted using MATLAB (MathWorks, v2018b) with *Smooth* function, using a smoothing parameter of 0.01.

### PDMS micropillars-array fabrication and fibronectin coating

Pillar fabrication was achieved by pouring PDMS, which was mixed at 10:1 with its curing agent (SYLGARD^TM^ 184, Dow Corning), over silicon molds with holes at fixed depths and geometric distance. The mold was then put upside down, onto a glass-bottom 35-mm dish (D35-20-0-N, Cellvis), which was followed by incubation at 60°C for 12 hours. After cooling to room temperature, the mold was peeled off while both the mold and the PDMS pillar cast were immersed in absolute ethanol to prevent micropillars from collapse. Afterwards, the ethanol was replaced by PBS, and then human plasma full-length fibronectin (FC010-10MG, Merck) was added to the dish, making a coating solution with final concentration of 10 μg/mL for a one-hour surface-coating at 37°C [32]. Finally, residual fibronectin was washed away by four serial replacements with serum-free Diamond’s TYI-S-33 medium. For the experiments using micropillars without fibronectin coating, absolute ethanol was replaced by PBS, which was then replaced by serum-free Diamond’s TYI-S-33 medium.

### Cell seeding on micropillar-arrays and time-series imaging

A total of 1 × 10^5 *E. histolytica* trophozoites was collected through tapping and subsequent centrifugation. These trophozoites were then seeded onto a fibronectin-coated (or non-coated) micropillar-array in a 35 mm glass-bottom dish filled with serum-free Diamond’s TYI-S-33 medium. The system was incubated in a heating chamber at 37 ℃ located on Zeiss LSM800 confocal microscope. Regarding *E. histolytica* trophozoites treated with bacteria, a concentration of 1 × 10^6 trophozoites/mL was incubated with *E. coli* O55 or *E. coli* K12 (1 × 10^9/mL) in serum-free Diamond’s TYI-S-33 medium at 37°C for 30 minutes while shaking. Afterwards, most bacteria were removed by washing the trophozoites with TYI medium through centrifugation at 726 × g for three cycles. The trophozoites were then resuspended in TYI medium, followed by seeding (1 × 10^5 trophozoites) onto a micropillar-array described above. Bright-field time-lapse imaging for *E. histolytica* on micropillars was performed using a 20× objective at 37°C. Images were scanned every four seconds with a time-series mode.

### Phalloidin staining on micropillars

A concentration of 1 × 10^6 trophozoites/mL was incubated with or without *E. coli* O55 or *E. coli* K12 (1 × 10^9/mL) in serum-free Diamond’s TYI-S-33 medium at 37°C for 30 minutes while shaking. Afterwards, most bacteria were removed by washing the trophozoites with TYI medium through centrifugation at 726 × g for three cycles. The trophozoites were then resuspended in TYI medium, followed by seeding onto a micropillar-array and incubated at 37°C for an hour. Then trophozoites were fixed by 4% paraformaldehyde (PFA) for 15 minutes at room temperature. After washing away PFA by PBS, the cells were permeabilized with 0.2% Triton-X for 10 minutes at room temperature. Afterwards, the cells were stained with Alexa Fluor^TM^ 488 Phalloidin (A12379, Invitrogen) with a dilution of 1:1000 for 15 minutes at room temperature. After another PBS wash, the cells were kept in PBS solution with SlowFade™ Gold Antifade Mountant (#S36937, Invertrogen), and then imaged by Zeiss LSM800 confocal microscope with a magnification of 20x with Airyscan processing.

Regarding three-dimensional reconstruction of F-actin structures, the cell is imaged with 63x objective scanned with Airyscan and Z-stack mode (0.2 µm interval between neighboring slices with a total thickness around 5 µm), followed by Airyscan processing and deconvolution.

### Velocity and directionality analysis

Cell motility analysis for velocity and directionality was performed with Fiji plugin *Manual Tracking* and software *Chemotaxis and Migration Tool* [73], using the same movies that were analyzed for micropillars displacements. Briefly, the position (x,y) coordinate of each cell was recorded by *Manual Tracking*. Then, the coordinates of the cells’ starting points are transformed to position (0,0) using *Chemotaxis and Migration Tool*, which is also used for calculating velocity and directionality of cells. Directionality is the ratio of the Euclidean distance to the accumulated distance between the starting point and the endpoint of a migrating cell. Trajectory plots are generated by R4.4.0 with *ggplot2*.

### Inhibitors treatment

To assess the effects of different inhibitors on *E. histolytica* traction force and migration, *E. histolytica* trophozoites were pre-incubated with E64d (10 µM for 16 hours), metronidazole (MNZ, 5 µM for 16 hours), or Cytochalasin D (CytD, 5 µM for 24 hours) at 37°C. Control trophozoites were treated with DMSO. A total of 1 × 10^5 trophozoites were collected via tapping and subsequent centrifugation. These trophozoites were then seeded onto a fibronectin-coated micropillar-array in a glass-bottom dish filled with serum-free Diamond’s TYI-S-33 medium, supplemented with the same concentration of inhibitors as used during pre-incubation, followed by live-imaging on a confocal microscope.

### Phalloidin staining on ibidi µ-Slide 8 Well plates

A total of 2 × 10^5 *E. histolytica t*rophozoites were collected via tapping and subsequent centrifugation. These trophozoites were then seeded onto an ibidi µ-Slide 8 Well plate (#80826, ibidi GmbH) pre-coated with or without fibronectin and incubated in serum-free Diamond’s TYI-S-33 medium at 37°C for one hour. Regarding *E. histolytica* trophozoites treated with bacteria, a concentration of 1 × 10^6 trophozoites/mL was incubated with *E. coli* O55 or *E. coli* K12 (1 × 10^9/mL) in serum-free Diamond’s TYI-S-33 medium at 37°C for 30 minutes while shaking. Afterwards, most bacteria were removed by washing the trophozoites with TYI medium through centrifugation at 726 × g for three cycles. The trophozoites were then resuspended in TYI medium, followed by seeding onto an ibidi plate and incubated at 37°C for one hour. Next, the plates were washed three times with PBS to remove residual TYI-S-33 medium. Trophozoites were then fixed with 4% PFA in PBS for 15 minutes at room temperature. Following PFA removal by PBS washing, cells were permeabilized with 0.2% Triton X-100 for 10 minutes at room temperature. Subsequently, the cells were stained with Alexa Fluor^TM^ 488 Phalloidin (A12379, Invitrogen) with a dilution of 1:1000 for 15 minutes at room temperature. Later, cells were washed with PBS and mounted with SlowFade™ Gold Antifade Mountant. Imaging was performed using confocal microscope with a 63x objective and Airyscan processing.

### Statistical analysis

Plots are generated by *R4.4.0* with *ggplot2* package with statistical analysis conducted utilizing Prism 9 (Graphpad Software Inc.). Unless otherwise specified, significance was assessed using one-way ANOVA for multiple comparisons (with Holm-Sidak method for *post-hoc* test) and unpaired two-tailed t-test for two groups of comparisons. Pearson correlation analysis and plots were produced by *R4.4.0* with *ggplot2* and *ggpubr* package. Chi-square test was performed by *gmodels* and *tableone* packages in R3.6.3.

## Acknowledgement

We would like to thank Joseph (Yossi) Klafter (Tel Aviv University) for helpful discussions on superdiffusive motion (Lévy-like Movement) and related analyses. Our sincere thanks also go to Malak Amer (Haguy Wolfenson lab) for support with the application of the micropillar-array system. Much appreciation goes to Fengming Hu (Faculty of Data and Decision Sciences, Technion) for discussions on the kinematics and mechanics of *E. histolytica* migration. We are also grateful to Lidan Shi (Haguy Wolfenson lab) for assistance with Fiji/MATLAB coding and support in experiments.

## Supporting information

**S1 Fig.**
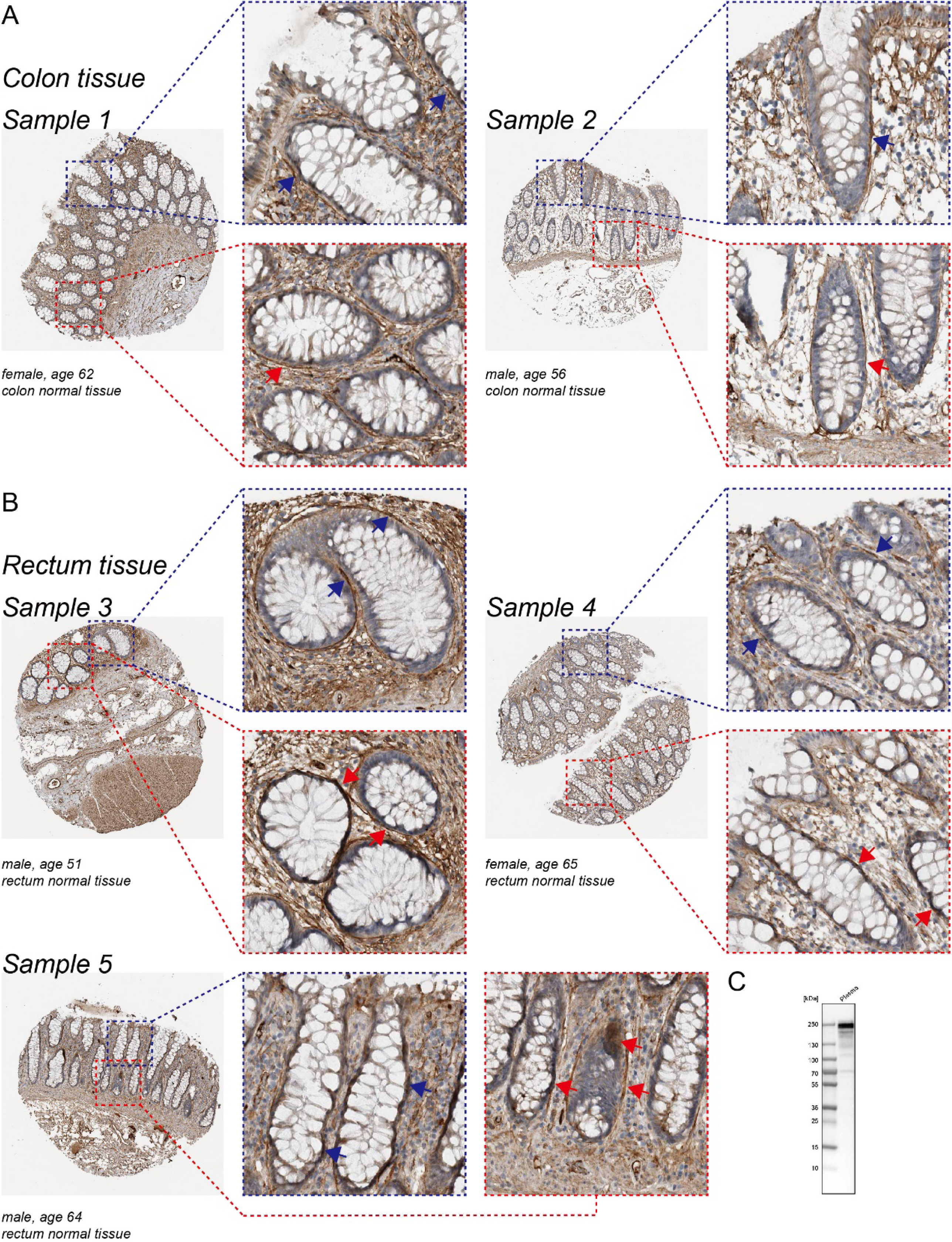
Fibronectin expression in human large intestinal mucosa. (A) Fibronectin expression in colon mucosa. (B) Fibronectin expression in rectum mucosa. Fibronectin is prominently expressed in the subepithelial basement membrane (indicated by arrows) and lamina propria. Images are sourced from the Human Protein Atlas (HPA) database (www.proteinatlas.org). (C) Western blotting results sourced from Atlas Antibodies (www.atlasantibodies.com) demonstrate the specificity of the antibody (HPA027066) used for staining the tissues shown in A and B.

**S2 Fig.**
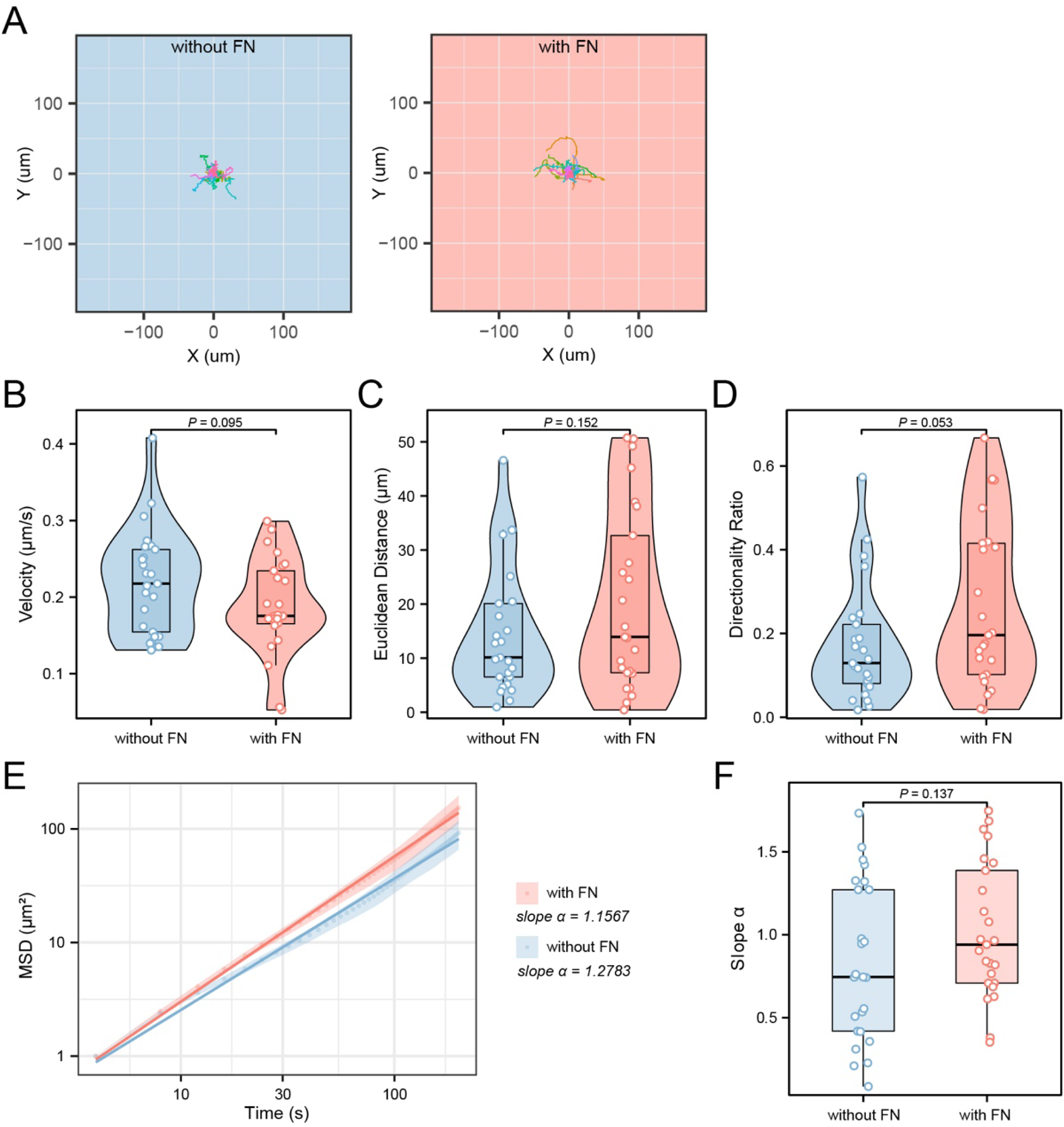
Motility properties of *E. histolytica* show no significant differences between fibronectin-coated and uncoated micropillars. (A) Trajectory plots of trophozoites on fibronectin (FN)-coated (right) and uncoated (left) micropillars. (B-D) Quantitative comparisons of velocity (B), Euclidean distance (C), and directionality (D) of *E. histolytica* between migrating on fibronectin-coated and uncoated micropillars. (E) The relationship between mean square displacements (MSD) and the corresponding time data was averaged and plotted on a logarithmic scale for *E. histolytica* locomotion under the different conditions. (F) Comparison of the slopes (α) for each individual cell/plot under the different conditions. For all the above diagrams, n = 25 in each group.

**S3 Fig.**
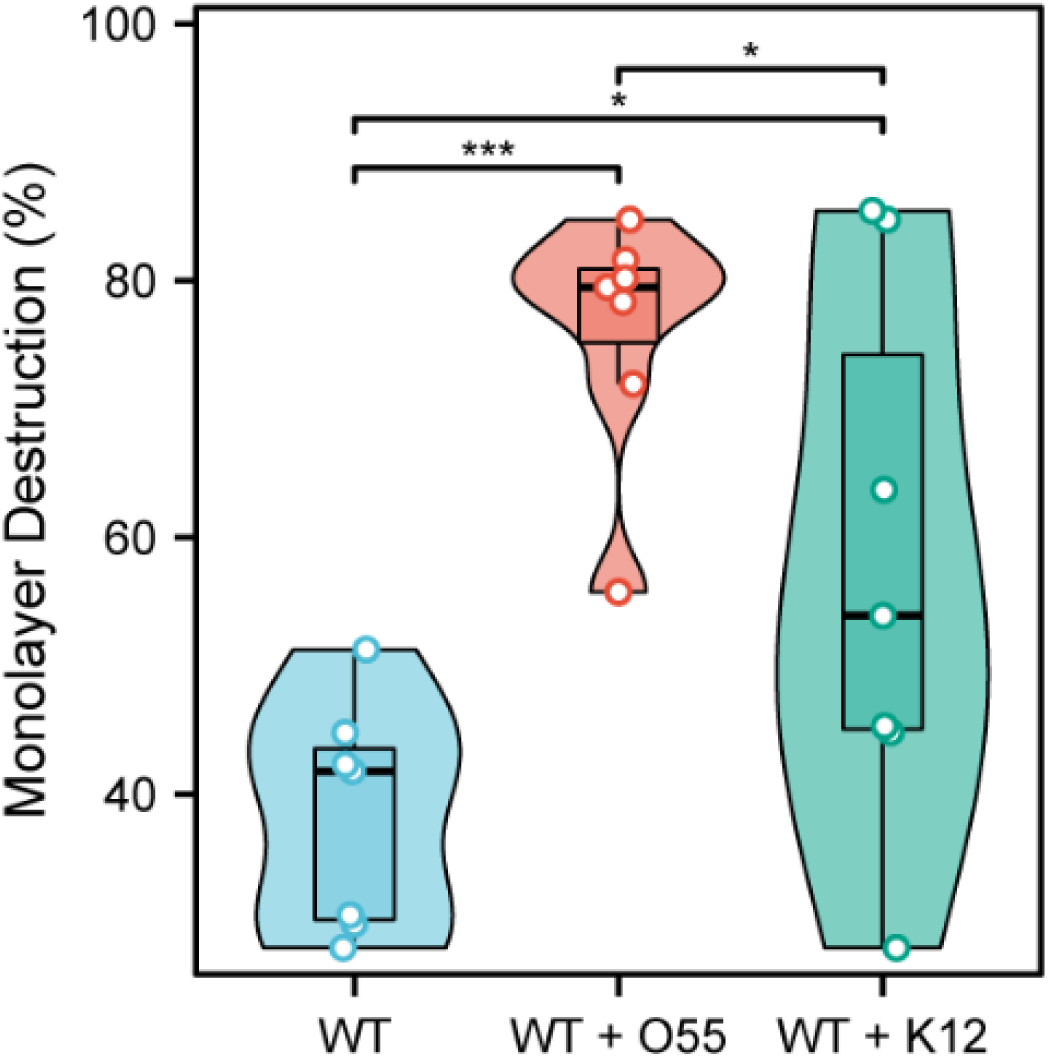
Cytopathic assay of *E. histolytica* trophozoites following short period of *E. coli* exposure. The cytopathic behavior of *E. histolytica* trophozoites, including controls (n = 7), trophozoites incubated with enteropathogenic *E. coli* O55 (n = 7), and trophozoites incubated with nonpathogenic *E. coli* K12 (n = 7), was assessed based on their ability to destroy a monolayer of HeLa cells. * *p* < 0.05, *** *P* < 0.001.

**S4 Fig.**
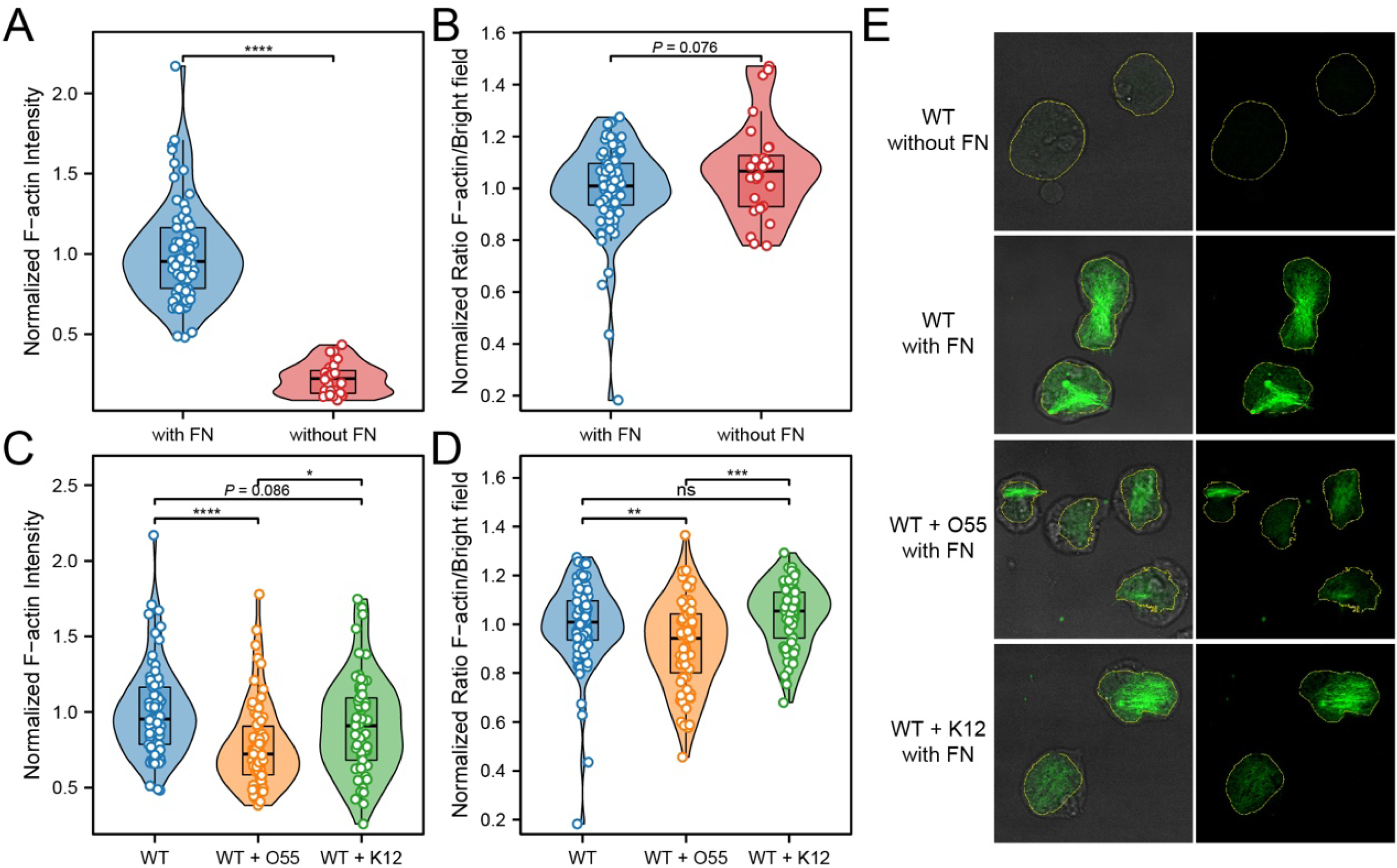
Alteration of F-actin morphology in *E. histolytica* migrating on ibidi µ-Slide 8 Well plates in different conditions. (A) Normalized Phalloidin intensity was quantitatively compared between *E. histolytica* trophozoites migrating on surface with (n = 77) and without (n = 28) fibronectin (FN) coating. (B) The normalized ratio of basal surface area (determined by confocal F-actin imaging) to the total area in the bright field (non-confocal imaging) was compared between trophozoites under two conditions of (A). (C) Phalloidin intensity was quantitatively compared among *E. histolytica* trophozoites under three conditions: control (n = 77), incubated with *E. coli* O55 (n = 67), and incubated with *E. coli* K12 (n = 61), on fibronectin (FN)-coated ibidi plates. (D) The normalized ratio of basal surface area (determined by confocal F-actin imaging) to the total area in the bright field (non-confocal imaging) was compared between trophozoites under three conditions of (C). (E) Representative images illustrate the differences in features compared in A-D. The yellow line outlines the basal surface area (determined by confocal F-actin imaging) of the cell. * *P* < 0.05, **** *P* < 0.0001.

**S1 Video. A representative video showing the motility of *E. histolytica* trophozoites on fibronectin-coated micropillar array, without exposure to *E. coli*.** Time stamp (minutes: seconds).

**S2 Video. A representative video showing the motility of *E. histolytica* trophozoites incubated with *E. coli* O55 on fibronectin-coated micropillar array.** Time stamp (minutes: seconds). In the videos, the curved organisms filling the screen are actually shadows of *E. coli*, distributed at different heights, due to the non-confocal nature of live imaging. In reality, the number of bacteria directly in contact with *E. histolytica* trophozoites is limited.

**S3 Video. A representative video cropped from Video 1 shows pillar displacement as an *E. histolytica* trophozoite migrates on a fibronectin-coated micropillar array, without *E. coli* exposure.** Time stamp (minutes: seconds). The trophozoite exhibits a non-directional exploratory behavior, actively using pseudopods to tug on the surrounding pillars. At the same time, the trophozoite seems to exhibit greater adhesion, resulting in more significant deflection of the micropillars.

**S4 Video. A representative video cropped from Video 2 shows pillar displacement as an *E. histolytica* trophozoite migrates on a fibronectin-coated micropillar array, incubated with *E. coli* O55.** Time stamp (minutes: seconds). When exposed to *E. coli* O55, the trophozoite transitions to rapid, goal-directed movement without actively pulling on the micropillars or engaging in random exploration, resulting in noticeably smaller micropillar deflections. In the videos, the curved organisms filling the screen are actually shadows of *E. coli*, distributed at different heights, due to the non- confocal nature of live imaging. In reality, the number of bacteria directly in contact with *E. histolytica* trophozoites is limited.

